# From Canopy Images to Organ-Level Disease Assessments: A Scalable Approach to Measure Quantitative Resistance in the Field

**DOI:** 10.1101/2025.04.30.651476

**Authors:** Radek Zenkl, Bruce A. McDonald, Achim Walter, Marie Unvericht, Cyrille Saintenac, Jonas Anderegg

## Abstract

Breeding for quantitative, polygenic resistance is widely considered the most durable, cost-effective, and environmentally safe approach to crop disease control. However, progress in resistance breeding is hindered by the limited capability of current approaches to measure highly quantitative disease phenotypes under field conditions with high precision and sufficient throughput.

Here, we present an imaging protocol and a modular image processing pipeline that enables wheat disease detection and severity estimation directly from very-high-resolution canopy imagery, eliminating the need for physical interaction with the monitored plants as required in previously proposed sensor-based methods capable of symptom-level diagnosis. The pipeline combines deep-learning-based semantic segmentation, keypoint detection, and depth estimation to diagnose and quantify disease symptoms and extract the analyzable reference plant surfaces for severity estimation. By leveraging estimated relative depth and analyzing image texture, well-focused areas with sufficient quality were accurately segmented. Despite the challenging nature of canopy images and frequent symptom ambiguity, symptom detection and segmentation models trained on a new dataset reached a similar performance as already described in more simplified scenarios where detached, flattened leaves were analyzed.

Plot-level severity estimates of Septoria Tritici Blotch, a major wheat disease, obtained using the new method and a precise but more laborious reference method were highly correlated (Pearson *R* = 0.83) across a range of morphologically contrasting cultivars. Validation of the new method on data collected by different operators at different sites demonstrated the robustness of the approach. The ability of the method to process imagery acquired in a contact-free manner can enable deployment on autonomous ground vehicles, paving the way for automated, scalable phenotype acquisition.

## 1 Introduction

Plant pathogens pose a significant threat to global food security, causing substantial yield and quality losses in major crops each year [58]. To mitigate these losses, modern agriculture relies primarily on two strategies: chemical control through application of pesticides and deployment of genetic resistance in host plants [23]. While the efficacy of pesticides is typically high, the widespread application of single-target-site pesticides exerts strong selection pressure on pathogens to evolve resistance [23], leading to a growing number of ineffective pesticides [28] whilst others were banned [44] due to associated collateral damage [66]. The development of new pesticides is slow and very expensive, leading to fewer pesticides being introduced [44].

Similar to pesticides, commercial germplasm typically contains a small number of resistance genes, also leading to high selection pressure [6] [42]. Cases of pathogens overcoming host resistance within a few years have been well documented [15] [57] [68] [13] [24] [37]. The mechanisms behind host resistance vary in terms of complexity and the degree of resistance they provide [53]. Multiple resistance mechanisms can be combined to increase effectiveness and durability of resistance by reducing selection pressure which can be imposed by single resistance mechanisms on pathogen populations [48] [51]. Breeding has primarily targeted single large-effect resistance genes because they are relatively easy to identify and introduce into commercial germplasm.

However, when exposed to diverse pathogen populations under field conditions, these large-effect resistance genes can still cause quantitative resistance phenotypes as a consequence of their varying effectiveness against different pathogen isolates [45] [46]. The combined effect of multiple such single-gene-mediated quantitative resistances within single cultivars can further diversify the overall resistance phenotypes observed in the field [9]. In addition, the presence of other types of resistance genes with smaller effects as well as escape mechanisms also contribute to the highly quantitative resistance reactions that evolve over time under field conditions.

Capturing these quantitative differences in resistance requires precise measurement of disease symptoms under field conditions [50] [1] [49]. If such measurements can be made at sufficient throughput, they can provide a new basis for identifying resistance genes with varying effect size and mechanisms, creating opportunities to combine genes or mechanisms for increased effectiveness and durability of host resistance [43].

In breeding programs, such quantification is still mostly conducted through expert visual scorings [8]. The associated need for trained personnel together with the subjective nature of visual scorings impose significant limitations in terms of throughput, availability, precision, and reliability of phenotypic data collected in this manner. This has led to a great interest in the development of potentially more objective and higher throughput sensor-based methods of disease assessment in the field. Previously proposed methods differ in the sensor types that they utilize and the scale that they operate on.

Approaches typically incorporate RGB or multi-spectral sensors among others (reviewed by [18]). Multi-spectral sensors offer increased contrast (e.g. between healthy and damaged tissue, [78]) and can reveal phenomena outside of the visible light spectrum (e.g. ultra-violet analysis, [11]). Hyperspectral sensors filter out a significant portion of the incoming light and thus require longer exposure times or an increase in sensor gain which amplifies sensor noise. With multi-lens multispectral cameras, depending on the imaging distance, individual channels can be difficult to map onto one another, as they observe the scene from a different viewing angle. Multispectral sensors with filter arrays capture each spectral band on separate pixel subsets, requiring spatial interpolation to reconstruct full images. On fine structures, this process can misalign edges between bands, creating artifacts such as blurring, color fringing, or false textures. Due to these limitations, RGB cameras may be the most suitable candidate for close-up imaging, particularly when considering variable and limited outdoor lighting.

Regardless of the sensor choice, existing approaches attempt to classify a disease [47] [21], localize affected plants or areas [36] [75], or even quantify the severity of the disease. The latter has been achieved through indirect estimation by image classification or regression of values from a pre-defined severity scale [26] [31] or by direct segmentation of the healthy and diseased areas [79] [25]. In this context, the term “disease detection” is often misused, as the phytopathological community typically refers to image classification (identifying which disease is present in an image) rather than true object detection, which also involves localization (pinpointing where the disease occurs in the image).

Deep learning has emerged as a dominant approach for processing visible symptoms on RGB images, leveraging both shape and color information, and requiring domain-specific datasets for effective training. While most of the developed methods introduce their own (typically small) dataset for one pathosystem to train models, an increasing number of larger datasets attempt to provide a broader basis for the development of more flexible approaches that can handle multiple pathosystems [30] [67] [75] [59] [63] [71] [72] [54]. While most of these datasets contain highly diverse data in the form of many diseases or locations, none distinguish more than one disease per image. As a result, models trained on them can perform diagnosis at the image level, which can be problematic because image background and symptom occurrence can be confounded. In contrast, when multiple diseases appear within a single image, diagnosis must occur at the symptom level, as the background does not offer reliable or decisive information for the classification of individual symptoms. These datasets can be grouped by imaging conditions (e.g. sampled leaves, uniform background, controlled lighting, in-field) and by the labels they provide (i.e., classification, detection, segmentation). Notably, the majority of the larger datasets [63] [72] [71] have been created by scraping data from the internet, using appropriate queries in contrast to sampling from designed experiments. Particularly for the case of in-field disease segmentation, the palette of available datasets remains limited [72] [54]. The PlantSeg dataset [72] lacks phenomena like senescence, chlorosis, and symptoms boundaries, limiting its usefulness for direct severity assessment. In contrast, the NLB dataset [54] covers one disease and one crop in-field but provides accurate boundaries of the disease symptoms. To the best of our knowledge, only the EFD dataset [79] combines outdoor lighting, co-infections within single images and high quality annotations suitable for symptom quantification and diagnosis. There is no dataset available capturing multiple diseases in images of field canopies.

Apart from detection, in-field disease quantification introduces additional challenges, as symptom quantification in an image must be combined with a measurement of an appropriate reference area analyzable within the same image. For some pathosystems, it may be sufficient to distinguish plants from the background when all visible parts of plants can be infected (e.g. Cercospora leaf spot in sugar beet). For others, it can be necessary to conduct the analysis on specific organs (e.g. Fusarium head blight in wheat). Fortunately, the scientific community recently laid significant ground work in the form of plant segmentation datasets [40] [12] [80] [5] [3] with the recent addition of a large and diverse organ segmentation dataset for wheat [70].

A typical example for a disease with highly quantitative resistance reactions that requires field evaluation is Septoria Tritici Blotch (STB), a major wheat disease caused by the fungus *Zymoseptoria tritici*. This disease causes visible symptoms on wheat leaves in the form of necrotic lesions and dark miniature fruiting bodies, called pycnidia. Pycnidia are typically used as a diagnostic feature of STB, as the necrotic lesions on their own cannot be easily distinguished from other diseases, stresses or senescence. STB progresses gradually, as epidemics are driven by secondary, asexual, reproduction cycles [81]. The number of pycnidia provides a proxy for reproductive potential and consequently potential disease pressure [52] [22].

STB severity is commonly estimated in the field using visual assessments of percentage of leaf area covered by lesions (PLACL) [56] [60] [20] [10]. However, this is not scalable and limits the volume of germplasm screened for breeding along with low repeatability due to inter-rater and intra-rater variance [8]. Furthermore, the resources required for each assessment result in low temporal resolution, as each assessment demands significant time and effort. Thus, many attempts for standardization, increasing the throughput and precision by utilization of optical sensors have been made. Previous works have either focused on necrotic lesions and omitted the recognition of pycnidia for the sake of in-field simple data acquisition [77] [2] [5] [7] [69], or considered the detection of pycnidia at the cost of tedious sample manipulation and simplified imaging conditions resulting in low throughput [4] [79] [64] [65] [41]. Notably, only [79] and [4] operated on a sufficient level of detail to distinguish different stressors and were able to robustly detect pycnidia under field conditions.

The aim of this work was to develop a standardized and scalable method for quantitative assessment of STB in wheat canopies. We developed a lean and readily transferrable imaging protocol that ensures sufficient image quality to enable non-invasive detection, diagnosis, and quantification of miniature disease symptoms directly in the field. The proposed modular, multi-step image processing pipeline quantifies symptoms and estimates plot disease severity directly from these canopy images. Its ability to capture canopy images from approximate viewpoints around plots makes it well-suited for deployment on autonomous machinery, which would facilitate scalable, automated disease severity assessments and potentially enable breeders to track disease progression over time and perform larger experiments.

## 2 Materials and Methods

### 2.1 Data Acquisition

As a basis for the development and validation of an image-based disease phenotyping method, a field experiment was carried out at the ETH Research Station for Plant Sciences in Lindau-Eschikon, Switzerland (47.449 N, 8.682 E) during the 2021-2022 and 2022-2023 wheat growing seasons. The main objective of this field experiment was to create broad variation among experimental plots in terms of visual canopy appearance and disease incidence and severity, similar to what would be encountered in a large breeding nursery. To this end, a set of morphologically diverse wheat cultivars was subjected to diverse treatments. Specifically, 16 cultivars were selected for their contrasting flag leaf angles, flag leaf glaucousness, and resistance to STB (for more details see Appendix A.2). Each cultivar was grown in multiple microplots sized 1.2m x 1.7m, which were subjected to the following treatments: i) an early fungicide application followed by artificial inoculation with 10 *Z. tritici* strains, ii) fungicide application without artificial inoculation, iii) neither fungicide application nor artificial inoculation (for more details see [5]).

Imaging was conducted using a full-frame mirrorless digital camera (EOS R5, Canon Inc., Tokyo, Japan; 45 megapixel, 36×24 mm sensor) combined with a macro lens (RF 35 mm f/1.8 IS Macro STM, Canon Inc., Tokyo, Japan). The aperture was set between F8.0 and F10.0 to balance depth of field, lens diffraction and exposure. Finding an optimal trade-off in this respect was essential because of the miniature size of pycnidia. Generally, the lens image is sharpest at aperture values of around F4.0. However, at such a narrow aperture, imaging objects from up close leads to the majority of the image being out of focus. Smaller apertures (i.e., higher F-numbers) increase depth-of-field but also introduce diffraction (interference of light on lens opening), which reduces overall image sharpness. Additionally, smaller apertures require longer exposure times, increasing the risk of motion blur. In our experiments, apertures between F8.0 and F10.0 provided the best operating point with a significant portion of the close-up image in focus while minimizing loss of sharpness throughout the entire image. The ISO was limited to 3200 to minimize sensor noise. The exposure time was kept below 1/60 s to minimize motion blur due to the operator’s movement or wind-induced motion of the plants. Both the camera and the lens are equipped with an active motion stabilization which mitigates the motion of the camera.

Image data was collected at various locations around a microplot from outside of the plot, pointing the camera at the border rows (see Appendix A.3). Locations were selected spontaneously and without reference to the presence and severity of diseases. The field of view of the camera was aligned horizontally so that flag leaves of the scanned canopy were central in the image. Thus, camera height was adjusted based on plant height rather than using one height for all plots. Although flag leaves were dominant in the resulting images, lower leaves could also be present due to in-plot variation in plant height, leading to vertical overlap between leaf layers. To ensure a consistent distance between the camera and the outermost parts of the canopy, an acrylic glass frame was attached to the lens of the camera and used as a gauge. The distance was approximately 18 cm, resulting in a sampling distance of 0.022 mm/px, comparable with 1200dpi (0.021 mm/px) scans from previous work ([79], [41], [64]). Compared to these works, our image acquisition protocol represents a step towards a more standardized and scalable way of imaging miniature features such as STB symptoms, favoring consistency across sites and institutions. Furthermore, this protocol only requires a consumer-grade camera and no extensive training of the operator.

To test the transferability of the imaging protocol and obtain an independent validation dataset from a different site, an operator from another research institution was trained and subsequently collected data independently, without on-site support. The training was completed within 30 minutes. This resulted in a secondary image dataset collected by INRAE in June 2024 in Landrethun-le-Nord (Pas-de-Calais, 50.854 N, 1.784 E), France, covering a bi-parental recombinant inbred line (RIL) mapping population from the two winter wheat cultivars ‘Winner’ (Florimond Desprez) and ‘Ultim’ (KWS) in generation F7. This population was chosen since the offspring showed a good variation in the degree of STB symptoms. The plots were exposed to naturally high disease pressure without artificial inoculation.

### 2.2 From Images to Deep Learning Datasets

Multiple datasets were created to support training and evaluation of deep learning models. All datasets were annotated at full resolution using the CVAT annotation tool [16]. A random 80/20% training-validation split of the newly annotated data from the application domain was used. Furthermore, when additional data from related works was available, it was incorporated as an extension of the training set, keeping the validation set identical. Such datasets were then named using a ‘+’ suffix to the original dataset name (e.g. EFD v2 and EFD v2+). For an overview of used datasets and their parameters see Table 1. For visual examples see Figure 1.

**Figure 1:**
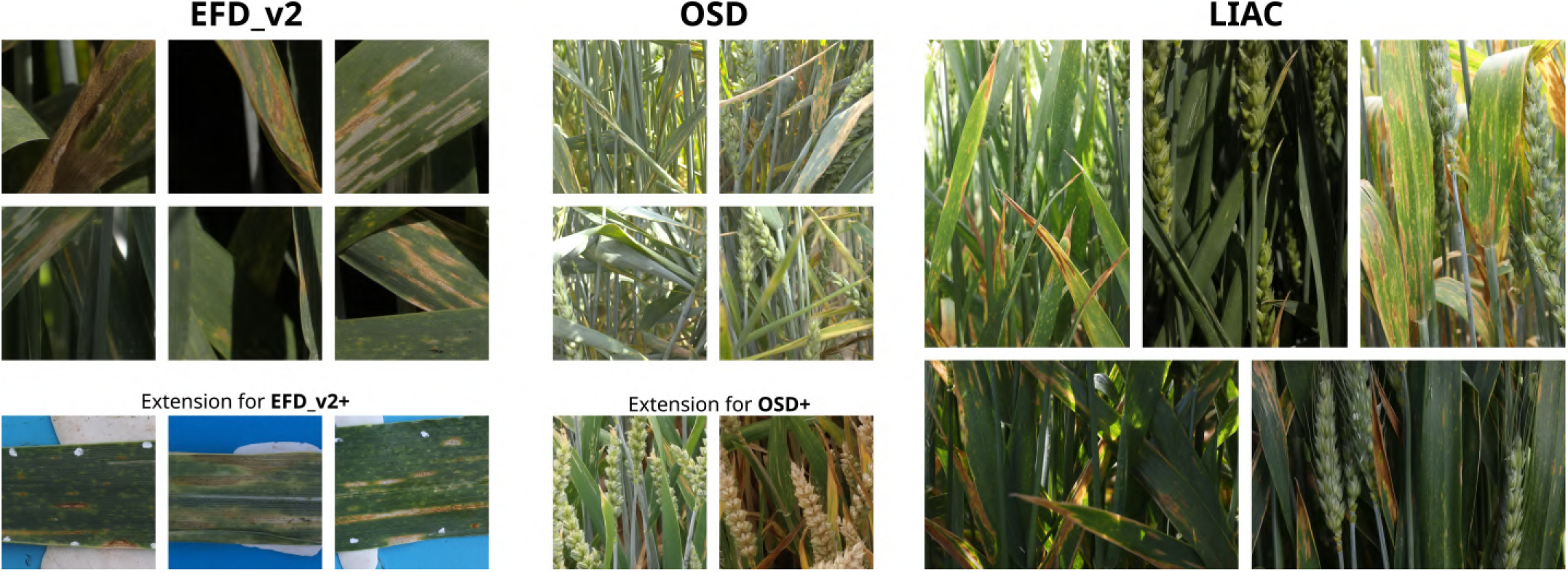
Example images from established deep learning datasets (EFD: Eschikon Foliar Disease dataset, OSD: Organ Segmentation Dataset, LIAC: Large unstructured images of canopy)

**Table 1:** Overview of manually annotated deep learning datasets (EFD: Eschikon Foliar Disease dataset, OSD: Organ Segmentation Dataset, LIAC: Large Images of Annotated Canopy)

| Dataset | # Images | Image Size | Annotated Classes | Years | Locations |
| --- | --- | --- | --- | --- | --- |
| OSD | 184 | 1024 x 1024 | Stems, Ears | 2023 | Eschikon |
| OSD+ | 321 | 1024 x 1024 | Stems, Ears |  | Eschikon |
| EFDv2 | 400 | 1024 x 1024 | Pycnidia, Rust Pustules, Necrosis, Insect Damage, Powdery Mildew | 2023 | Eschikon |
| EFDv2+ | 1081 | 1024 x 1024 | Pycnidia, Rust Pustules, Necrosis, Insect Damage, Powdery Mildew | 2022, 2023 | Eschikon |
| LIAC | 40 | 6144 x 4096,<br>4096 x 6144 | Stems, Ears, Pycnidia, Rust Pustules, Necrosis, Insect Damage, Powdery Mildew | 2023, 2024 | Eschikon,<br>Landrethun-le-Nord |

An accurate recognition of disease symptoms is the main prerequisite for any downstream analysis of disease-related phenotypes. As a first step towards this goal, we compiled a dataset providing annotated disease symptoms for model development. Following the concept of the previously established leaf-level EFD dataset [79], we annotated 400 image crops of unstructured canopy (see Section 3.1 with size 1024 x 1024 px. The regions were selected manually on randomly sampled images to ensure presence of disease symptoms at sufficient image quality (see Figure 1). However, due to the unstructured nature of canopy images, most of the resulting image crops also contained regions exhibiting low quality, typically due to focus blur. Symptoms were annotated as long as they could be diagnosed with high confidence, often leading to annotations of necrosis or rust pustules in substantially blurred regions. In contrast to our previous work, we omitted the leaf class given implicit extraction of the leaf area as a result of organ segmentation and focus estimation (see Section 3.3). In the newly sampled images for annotation, we discovered that the presence of powdery mildew (caused by *Blumeria graminis* f. sp. *tritici*) led to false positives for lesions and pycnidia, probably because powdery mildew fruiting bodies (chasmothecia) are very similar in shape, size and color to STB pycnidia. To explicitly consider this scenario during training and evaluation, powdery mildew was annotated with polygons. This formed the basis for a new EFDv2 dataset which features symptom-level annotation in a natural environment, adding variability from scene composition. Compared to existing leaf-level datasets, these images include extra plant organs, diverse lighting conditions, and changing object orientations, affecting image quality through focus blur and viewing angle shifts.

Yet, the previously annotated images of flattened leaves may still be useful as the tasks of symptom detection and segmentation are likely highly correlated due to the consistent appearance of symptoms regardless of leaf orientation. To test this hypothesis, we compiled the EFDv2+ dataset from the newly generated canopy-level and pre-existing leaf-level annotated images ([4], [79]). By keeping the validation sets identical, a direct comparison and quantification of the impact of including samples of flattened leaves can be made.

Since objects are not pre-selected (e.g. through leaf sampling), the method must identify the relevant areas for evaluation. For plant diseases, this means identifying colonizable organs (e.g., leaves for STB). To support this, we created the Organ Segmentation Dataset (OSD) with 184 randomly selected, 4096×4096px cropped images (see Figure 1), annotated for stems and ears. Awns were not annotated, and blurred objects were marked as background.

The proposed image processing pipeline (see Section 3.3) extracts the relevant leaf regions implicitly, without the need for explicit leaf labels. For model development, images and labels were down-scaled by a factor of 4 to 1024 x 1024 px to reduce the computational complexity. The resulting OSD training set was complemented with 137 annotated images from the same site but collected using a different imaging setup [3] to compile an OSD+ dataset.

Finally, to validate the complete evaluation of diseases from canopy images end-to-end, we compiled a Large Images of Annotated Canopy dataset (LIAC). This dataset consists of 20 images from Eschikon and 20 images from Landrethun-le-Nord (see Section 3.1) taken at both portrait and landscape orientations and cropped to 6144 x 4096 px (see Figure 1). These images were annotated with both the disease symptoms labels and organs labels as described above. The goal of the LIAC dataset is to enable a direct comparison with respect to a human without any intermediate steps. Due to two different locations, this dataset should also provide insights into the transferability of the acquisition protocol and performance of the developed methodology when used on data originating from a previously unseen location.

### 2.3 Image Processing Method

Our method builds on [79] and enhances its ability to analyze foliar diseases in unstructured canopy images. It retains the original symptom detection and segmentation approach but adds two stages for focus segmentation and plant organ segmentation to refine the regions for symptom analysis. Specifically, the method employs four deep learning models with distinct semantic tasks and scales (see Figure 2).

**Figure 2:**
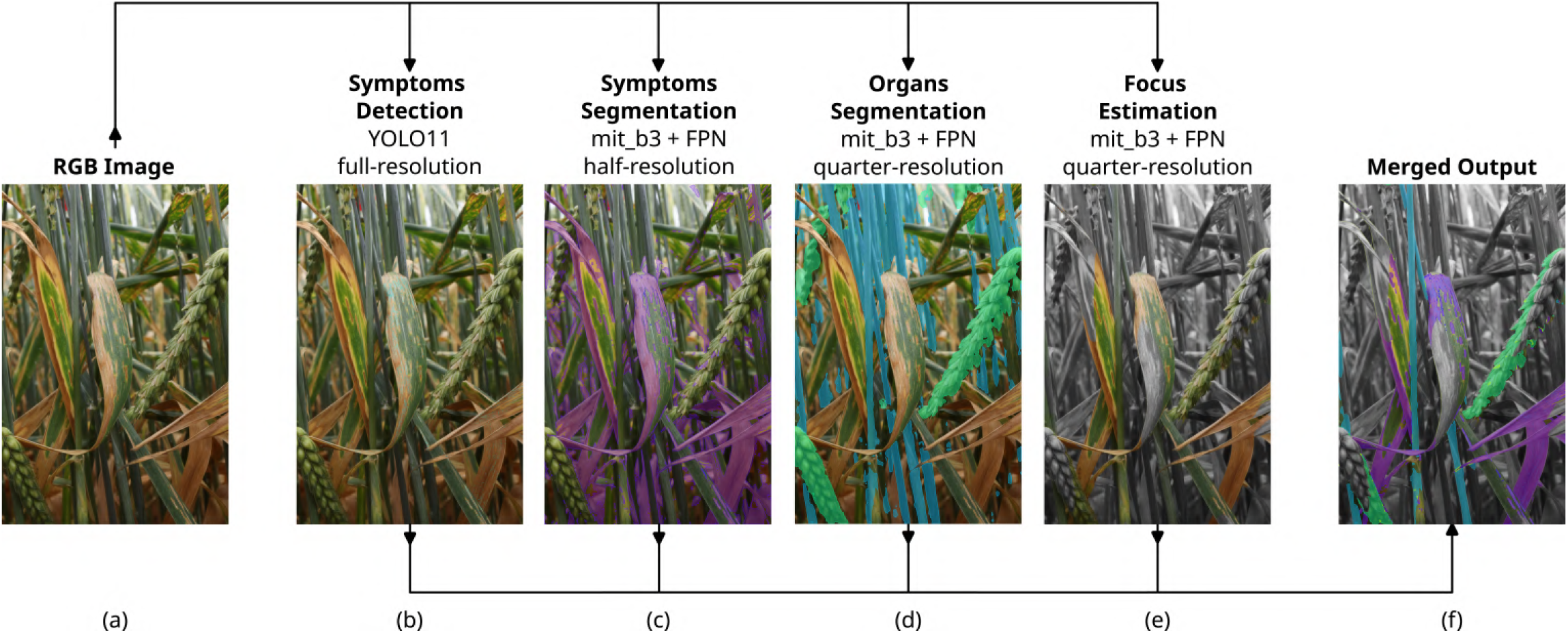
Overview of the image processing pipeline.

The first building block is detecting STB pycnidia and rust pustules, based on YOLOv11 Pose model [33], the successor architecture of YOLOv8 [32] used in previous work [79]. Next, symptoms segmentation model consists of Seg-former backbone [74] and FPN head [35]. Compared to previous work [79] the data augmentation of both keypoint detection and symptoms segmentation models was adjusted to facilitate for random rotations, as the orientation of the symptomatic leaves in a canopy can vary within one image or across different images. Furthermore, to allow for processing of large images without the need for splitting into patches, the input for the symptoms segmentation model was downscaled by a factor of 2 to reduce the memory requirements during inference. The next building block is organ segmentation. We employed the same architecture as for the symptoms segmentation, i.e., Segformer backbone [74] and FPN head [35]. By segmenting leaves and ears, leaves are extracted implicitly as the remainder.

The last step executes focus estimation where a novel transformer based Depth Anything v2 [76] model predicts relative depth from a monocular RGB image which is combined with classical computer vision analysis of texture. For the latter, high-frequency features, characterized by abrupt changes in image intensity, were detected. These features should only appear in regions that are in focus, where fine details and sharp edges are preserved. Conversely, blurred regions predominantly exhibit low-frequency characteristics, meaning intensity changes occur more gradually. This is due to the smoothing effect introduced by defocus, which reduces sharp transitions and blends adjacent areas together. To detect high-frequency features we used a thresholded difference of a Gaussian (DoG) in form of a convolution with a discrete gaussian kernel of size of 5 and 11. The kernel sizes were determined by an approximation of the size of pycnidia in the image so that features of similar size are extracted by the filter. The DoG was computed separately for each color channel and features occurring in all three channels were accepted, since we expect fine texture to lead to changes in overall intensity rather than a change in a singular color channel. To reject images which are completely blurry, at least 10’000 such features are required. Finally, the predicted depth from Depth Anything v2 at which the most features appear is chosen as the estimated focal plane (see dashed line in Figure 3d). The in-focus depth range is then determined based on the statistical distribution of detected features, as absolute depth values are not available. To refine this estimate, outliers beyond 3 ∗ *σ* from the mean are removed from the depth histogram. The width of the in-focus depth range is then set based on the variance of the remaining depth values. For this we used the formula: *λ* ∗ *σ*, where lambda is a tunable parameter which controls the aggressiveness of the focus estimation.

**Figure 3:**
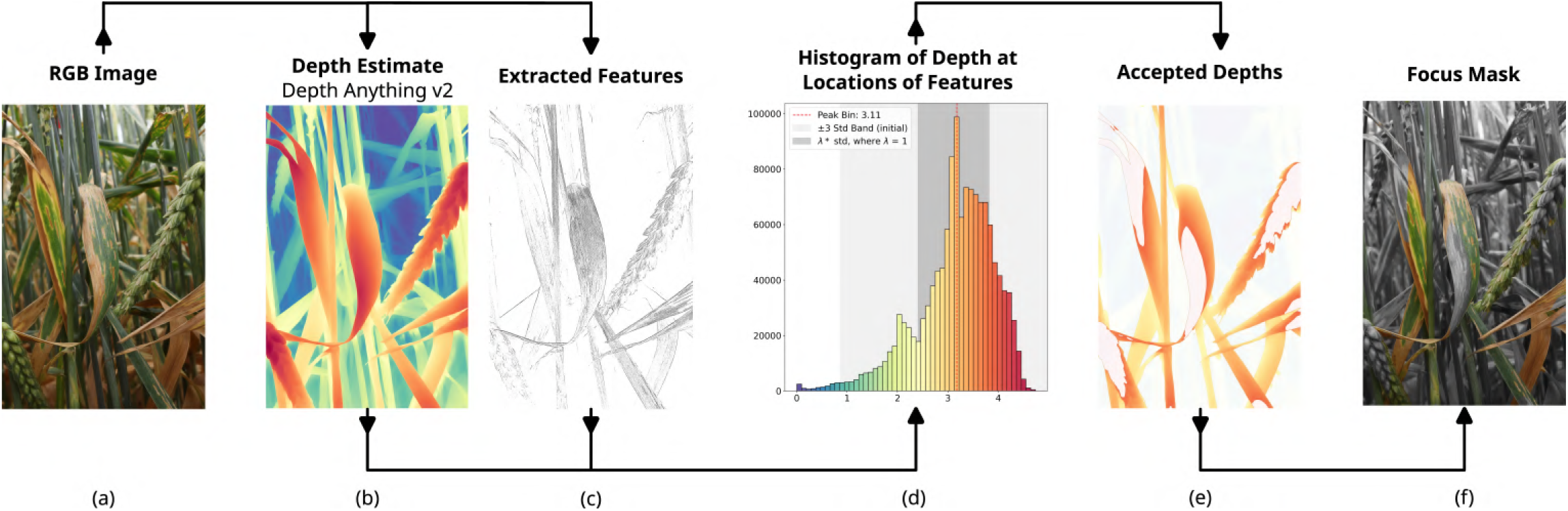
Overview of the focus estimate pipeline.

This approach allows us to use sparse feature detection (see Figure 3c) to identify sharp regions while incorporating smooth depth predictions from Depth Anything v2 (see Figure 3b)to define clear and gradual boundaries around infocus areas (see Figure 3f). Both the Depth Anything v2 model and DoG were applied on images scaled down by a factor of 4 to reduce computational complexity and suppress sensor noise.

All the described modules run in parallel and their results are saved independently to allow for processing large batches of data step by step. Leveraging the combined output of the organ segmentation and focus estimation blocks described above, we are capable of understanding the scene and subdividing it into different semantic areas. In the context of STB, we can extract the relevant area, i.e. leaf surfaces in focus, and use this as the region of interest for the downstream symptom analysis.

### 2.4 Hyperparameter Optimization

To optimize the model performance, we tuned model hyperparameters within a limited search space. Specifically, we explored different backbone depths and varying learning rate, momentum and weight decay for the AdamW optimizer [39] repeated on all corresponding datasets (see Table 1).

Even though the same architecture is used for both organs segmentation and symptoms segmentation, they were optimized and trained separately to retain the individual blocks’ modularity and independency. Since the goal of this work is to process large images similar to the LIAC dataset (see Table 1), the hardware requirements and runtime needs to be considered. The segmentation block is the computationally most expensive part of the complete pipeline due to Segformer’s attention mechanism which runs O(*n*^2^) which potentially requires the largest amount of memory. For organ segmentation, the computational cost was mitigated by down-scaling the input data by factor 4 since the additional resolution did not yield any benefit to the performance. However, for symptoms segmentation the full resolution can be beneficial during the analysis. Thus, different scaling factors of 1 (native resolution), 1/2 and 1/4 were tested. These scaling factors were selected so that the resolution is down-scaled homogeneously in the whole image as the down-scaling grid is always aligned with the original image, thus decreasing the risk of creating scaling artifacts.

For inference, the available GPU Memory is a factor to be considered. In the scope of this work, 24 Gb of VRAM per model were available. When a single inference sample did not fit into the GPU memory as a whole, it needed to be subdivided into patches which were processed individually. This can introduce an unintended drop in performance due to the decreased receptive field and increased amount of image borders which typically exhibit worse performance.

### 2.5 Focus Estimation Tuning

The proposed approach for focus estimation allows for tuning the cutoff depth for in-focus region prediction by tuning the parameter *λ* (see Section 3.3). The parameter *λ* directly drives the aggressiveness of the focus estimation, leading to larger or smaller fractions of the image being accepted as in-focus. We expected more aggressive focus estimation to lead to higher accuracy of disease recognition, as only image regions of the highest quality are evaluated. In contrast, less aggressive focus estimation yields a higher fraction of the image for evaluation. Ideally, focus estimation would be tuned in such a way so that the largest potential throughput is achieved without a significant decline of accuracy.

To analyze this behavior, the LIAC dataset (see Table 1) was processed multiple times with the exact same pipeline, only changing the focus estimation aggressiveness. This created a basis for assessing the accuracy of pycnidia detection and necrosis segmentation within the regions predicted to be in focus. For this, pycnidia F1 score and necrosis IoU were monitored while the focus estimation parameter was modified. In this analysis, a detected keypoint was considered correct when within 5 px of the annotated counterpart. Furthermore, to suppress the influence of samples with low presence of one particular class, for necrosis at least 1% had to be present, for organs at least 5% had to be present and for pycnidia and rust pustules at least 25 instances had to be present. The filtering was conducted for the corresponding class only.

### 2.6 Proof of Concept

While annotated datasets are valuable for evaluating performance on individual images, they offer limited insight into estimating values at the plot level. To assess performance on individual plots, further images were collected on 12.6.2023, in accordance to the proposed image acquisition protocol. At each of the 71 microplots with an STB incidence of at least 33%, images were captured from eight different locations around the plot. Each imaging location was evaluated individually with plot values estimated by computing a weighted average, with weights corresponding to the size of the estimated reference area (see Section 3.3). In parallel to the imaging campaigns, reference data on disease incidence and severity was collected on 15.6.2023. The purpose of the reference method was to obtain as objective and precise an estimate of STB severity as possible while minimizing the potential scorer bias. Following a protocol described earlier [2], disease incidence on flag leaves was visually assessed for 30 culms per plot and then multiplied by a conditional severity measure quantifying the percentage of leaf area covered by lesions obtained from eight detached and scanned infected flag leaves per plot. To select the leaves, a random culm was chosen without prior knowledge of symptom presence. If no STB symptoms were observed, a new culm was inspected. If symptoms were present, the flag leaf was detached and imaged using a flatbed scanner, following the method described by Stewart et al. [64]. The resulting reference values yielded the opportunity to put the proposed method into perspective with an established methodology, where extracted traits in the form of mean PLACL and pycnidia density plot values can be directly compared.

## 3 Results

### 3.1 Model Performance

Overall for symptom segmentation, all models performed best in segmenting necrosis, followed by insect damage, with the lowest performance observed for powdery mildew (Table 2). This trend reflects the proportional representation of these classes in the datasets, as lower label proportions generally corresponded to reduced model performance.

**Table 2:** Comparison of performance for different backbones of SegFormer trained on multiple datasets for symptoms segmentation: EFD [79] - flattened leaves; EFDv2 - canopy images; EFD v2+ - flattened leaves and canopy images. Values for IoU and F1 score for necrosis, insect damage, and powdery mildew are reported. Note that EFD dataset does not contain labels for powdery mildew.

| Backbone | Dataset | Necrosis |  | Insect Damage |  | Powdery Mildew |  |
| --- | --- | --- | --- | --- | --- | --- | --- |
|  |  | IoU | F1 | IoU | F1 | IoU | F1 |
| mit_b0 | EFD | 0.59413 | 0.74540 | 0.2130 | 0.35120 | N/A | N/A |
| mit_b1 |  | 0.58311 | 0.73667 | 0.1578 | 0.27258 | N/A | N/A |
| mit_b2 |  | 0.54630 | 0.70659 | 0.17209 | 0.29364 | N/A | N/A |
| mit_b3 |  | 0.61628 | 0.76259 | 0.24903 | 0.39876 | N/A | N/A |
| mit_b4 |  | 0.61454 | 0.76125 | 0.19564 | 0.32726 | N/A | N/A |
| mit_b0 | EFD_v2 | 0.71208 | 0.83183 | 0.59818 | 0.74858 | 0.26105 | 0.41402 |
| mit_b1 |  | 0.72710 | 0.84199 | **0.62167** | **0.7667** | 0.34949 | 0.51795 |
| mit_b2 |  | 0.72407 | 0.83995 | 0.57940 | 0.73370 | 0.35315 | 0.52197 |
| mit_b3 |  | **0.73334** | **0.84616** | 0.60264 | 0.75206 | **0.45630** | **0.62666** |
| mit_b4 |  | 0.72027 | 0.83739 | 0.58386 | 0.73726 | 0.30313 | 0.46523 |
| mit_b0 | EFD_v2+ | 0.71302 | 0.83247 | 0.61839 | 0.76420 | 0.11603 | 0.20794 |
| mit_b1 |  | 0.72001 | 0.83721 | 0.59399 | 0.74529 | 0.30741 | 0.47026 |
| mit_b2 |  | 0.71997 | 0.83719 | 0.60370 | 0.75288 | 0.32382 | 0.48922 |
| mit_b3 |  | 0.73197 | 0.84525 | 0.61247 | 0.75967 | 0.39196 | 0.56318 |
| mit_b4 |  | 0.71424 | 0.83330 | 0.56856 | 0.72494 | 0.29669 | 0.45761 |

Deeper backbones generally outperformed shallower ones. However, these improvements were subtle compared to performance fluctuations inherent to the training process. Among all tested datasets, the encoder mit b3 consistently achieved the highest performance (see mit b3 in Table 2).

The segmentation model’s performance varied significantly depending on the training dataset. The greatest improvement was observed when training data originated from the same acquisition method as the validation set (see Table 2). This underscores the domain gap between flattened leaves imaged according to [79] and those obtained via the proposed free-form imaging. The largest performance difference was observed for the insect damage class, likely due to the characteristic blue background visible through damaged leaf tissue in the EFD dataset, which was not present in EFDv2 data.

Segmentation models exhibited comparable performance for necrosis and insect damage across the EFD v2 and EFD v2+ datasets. However, a significant gap was observed in powdery mildew segmentation, where models trained on EFD v2 consistently outperformed those trained on EFD v2+ (see Table 2). Furthermore, despite EFD v2+ being more than twice the size of EFD v2 (see Table 1), no significant improvement in symptom segmentation performance was observed.

For keypoint detection, both deeper models and larger datasets generally improved performance (Table 3). However, this trend did not always hold for the optimal confidence threshold, as better performance in terms of F1 score or mAP(50) did not necessarily yield higher confidence values (see Table 3). The most notable performance boost was observed when training data originated from the same domain as the validation set, specifically datasets EFD v2 and EFD v2+.

**Table 3:** Comparison of performance for different backbones of Yolo11 trained on multiple datasets for symptoms segmentation: EFD [79] - flattened leaves; EFDv2 - canopy images; EFD v2+ - flattened leaves and canopy images. Values for mAP(50), F1 score and confidence are report, where confidence refers to the optimal confidence threshold value to achieve the maximal F1 score for a given class.

| Model | Dataset | Pycnidia |  |  | Rust |  |  |
| --- | --- | --- | --- | --- | --- | --- | --- |
|  |  | mAP(50) | F1 | Confidence | mAP(50) | F1 | Confidence |
| Yolo11n-pose | EFD | 0.488 | 0.486 | 0.030 | 0.514 | 0.511 | 0.071 |
| Yolo11s-pose |  | 0.513 | 0.503 | 0.030 | 0.531 | 0.522 | 0.071 |
| Yolo11m-pose |  | 0.548 | 0.535 | 0.030 | 0.559 | 0.539 | 0.071 |
| Yolo11l-pose |  | 0.542 | 0.528 | 0.030 | 0.526 | 0.527 | 0.081 |
| Yolo11n-pose | EFD_v2 | 0.594 | 0.589 | 0.172 | 0.647 | 0.624 | 0.273 |
| Yolo11s-pose |  | 0.643 | 0.631 | 0.212 | 0.653 | 0.635 | 0.333 |
| Yolo11m-pose |  | 0.658 | 0.640 | 0.152 | 0.689 | 0.663 | 0.253 |
| Yolo11l-pose |  | 0.653 | 0.636 | 0.162 | 0.709 | 0.669 | 0.222 |
| Yolo11n-pose | EFD_v2+ | 0.607 | 0.598 | 0.172 | 0.713 | 0.673 | 0.242 |
| Yolo11s-pose |  | 0.651 | 0.639 | **0.263** | 0.740 | 0.690 | 0.253 |
| Yolo11m-pose |  | 0.653 | 0.635 | 0.202 | 0.736 | 0.692 | **0.374** |
| Yolo11l-pose |  | **0.665** | **0.650** | 0.212 | **0.741** | **0.698** | 0.283 |

In contrast to the segmentation models, the best detection performance was achieved with the EFD v2+ dataset. This might be linked to the fact, that visual appearance of rust pustules and pycnidia is less influenced by the change of imaging methodology (i.e., going from flattened leaves to canopy imaging) which makes it easier to use data from both scenarios. The reported confidence values correspond to the optimal confidence threshold for each class, maximizing the F1 score (i.e. the geometric mean of precision and recall). While lower confidence values were typically associated with poorer performance (as seen with the EFD dataset in Table 3), the highest performance did not always correspond to the highest confidence scores.

With the exception of models trained on the EFD dataset, the detection of rust pustules consistently outperformed the detection of pycnidia.

For organ segmentation, the performance on ears was consistently higher than for stems (Table 4). Interestingly, the additional training data contained in OSD+ did not consistently improve segmentation results across all classes. This is particularly evident for stems, where the highest IoU and F1 scores were achieved with OSD, indicating that domain differences in OSD+ may have introduced variability rather than a performance gain.

**Table 4:** Comparison of performance for different backbones of SegFormer trained two different datasets for organ segmentation: OSD - annotated images collected with the proposed acquision protocol; OSD+ - extension of OSD by including annotated canopy images using a different acquisition protocol [3]. Values for IoU and F1 score for ears and stems are reported.

| Backbone | Dataset | Ears |  | Stems |  |
| --- | --- | --- | --- | --- | --- |
|  |  | IoU | F1 | IoU | F1 |
| mit_b0 | OSD | 0.7954 | 0.88622 | 0.64877 | 0.78697 |
| mit_b1 |  | 0.82264 | 0.90269 | 0.67564 | 0.80643 |
| mit_b2 |  | 0.82555 | 0.90444 | 0.66415 | 0.79818 |
| mit_b3 |  | 0.81917 | 0.90060 | **0.67891** | **0.80875** |
| mit_b0 | OSD+ | 0.79744 | 0.88731 | 0.62946 | 0.7726 |
| mit_b1 |  | 0.79141 | 0.88156 | 0.62907 | 0.77122 |
| mit_b2 |  | **0.82955** | **0.90684** | 0.67583 | 0.80656 |
| mit_b3 |  | 0.82087 | 0.90163 | 0.64930 | 0.78736 |

For symptoms segmentation, using higher resolution generally led to better performance (Table 5). Mit b3 achieved the highest performance at all input sizes. Moving from quarter-resolution to full-resolution improved the performance substantially across all tracked metrics. The difference between half-resolution and full-resolution was smaller and less conclusive, as mit b3 with half-resolution achieved a higher score for powdery mildew than with full-resolution. However, for all other classes, increasing the input size consistently improved the performance across all backbones.

**Table 5:** Comparison of performance of SegFormer utilizing different input sizes for the EFDv2+ dataset.

| Backbone | Resolution | Necrosis |  | Insect Damage |  | Powdery Mildew |  |
| --- | --- | --- | --- | --- | --- | --- | --- |
|  |  | IoU | F1 | IoU | F1 | IoU | F1 |
| mit_b0 | quarter-resolution<br>(256 × 256 px) | 0.66353 | 0.79774 | 0.54293 | 0.70377 | 0.089485 | 0.16427 |
| mit_b1 |  | 0.68329 | 0.81185 | 0.55044 | 0.71004 | 0.080017 | 0.14818 |
| mit_b2 |  | 0.67167 | 0.80359 | 0.53724 | 0.69897 | 0.17017 | 0.29085 |
| mit_b3 |  | 0.69156 | 0.81766 | 0.55879 | 0.71696 | 0.13572 | 0.23900 |
| mit_b0 | half-resolution<br>(512 × 512 px) | 0.71302 | 0.83247 | 0.61839 | 0.76420 | 0.11603 | 0.20794 |
| mit_b1 |  | 0.72001 | 0.83721 | 0.59399 | 0.74529 | 0.30741 | 0.47026 |
| mit_b2 |  | 0.71997 | 0.83719 | 0.60370 | 0.75288 | 0.32382 | 0.48922 |
| mit_b3 |  | 0.73197 | 0.84525 | 0.61247 | 0.75967 | **0.39196** | **0.56318** |
| mit_b0 | full-resolution<br>(1024 × 1024 px) | 0.73098 | 0.84459 | 0.60923 | 0.7517 | 0.20527 | 0.34063 |
| mit_b1 |  | 0.73279 | 0.84579 | 0.61350 | 0.76046 | 0.35374 | 0.52261 |
| mit_b2 |  | 0.73545 | 0.84756 | 0.64529 | 0.78441 | 0.34357 | 0.51143 |
| mit_b3 |  | **0.74017** | **0.85069** | **0.64738** | **0.78595** | 0.35521 | 0.52421 |

### 3.2 Pipeline Benchmark

To test the performance of the overall pipeline end-to-end (i.e., without any prior region selection done by humans), the segmentation and keypoint detection models were applied on the LIAC dataset, originating from two separate field experiments (see Table 1 and Figure 1). Overall, the F1-score of pycnidia detection is comparable with the values reported on the validation set, but the detection of rust pustules achieved worse performance then on the validation set (see Figure 4 and Table 3). In terms of organ segmentation, there was a consistent drop of performance for both ears and stems (see Figure 4 and Table 4).

**Figure 4:**
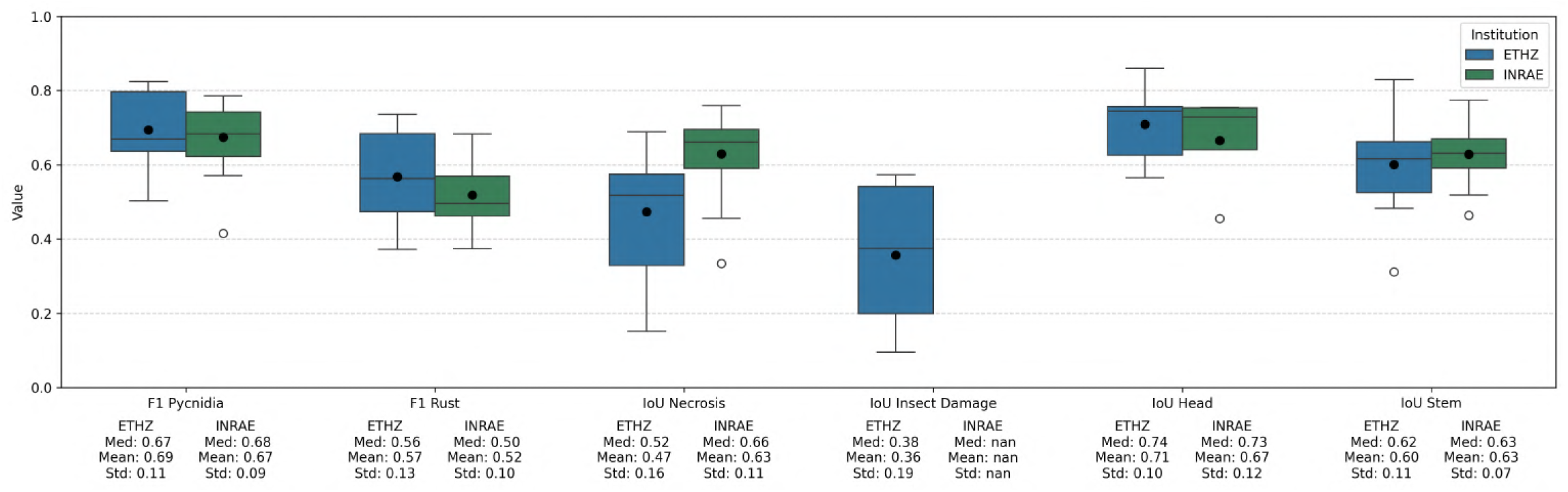
Boxplot of key metrics for keypoint detection, symptoms segmentation and organ segmentation on the LIAC dataset consisting of large unstructured images of canopy. For keypoint detection F1 score is reported and for segmentation IoU is reported. Metrics are plotted separately for the two subsets (ETHZ and INRAE). Note that insect damage on INRAE samples was not sufficiently prevalent to be numerically evaluated and compared with ETHZ.

Mean and median performance of keypoint detection and organ segmentation was comparable across experiments (Figure 4). However, symptom segmentation performance was higher at INRAE than at ETHZ subset.

The impact of a focus estimation aggressiveness on necrosis IoU and pycnidia F1 score is depicted in Figure 5. For the segmentation of necrosis, there was a clear trend, where more aggressive focus estimation led to higher performance. Surprisingly, the effect was negligible for the performance of pycnidia detection, which appeared to be insensitive to the selection of focus parameter.

**Figure 5:**
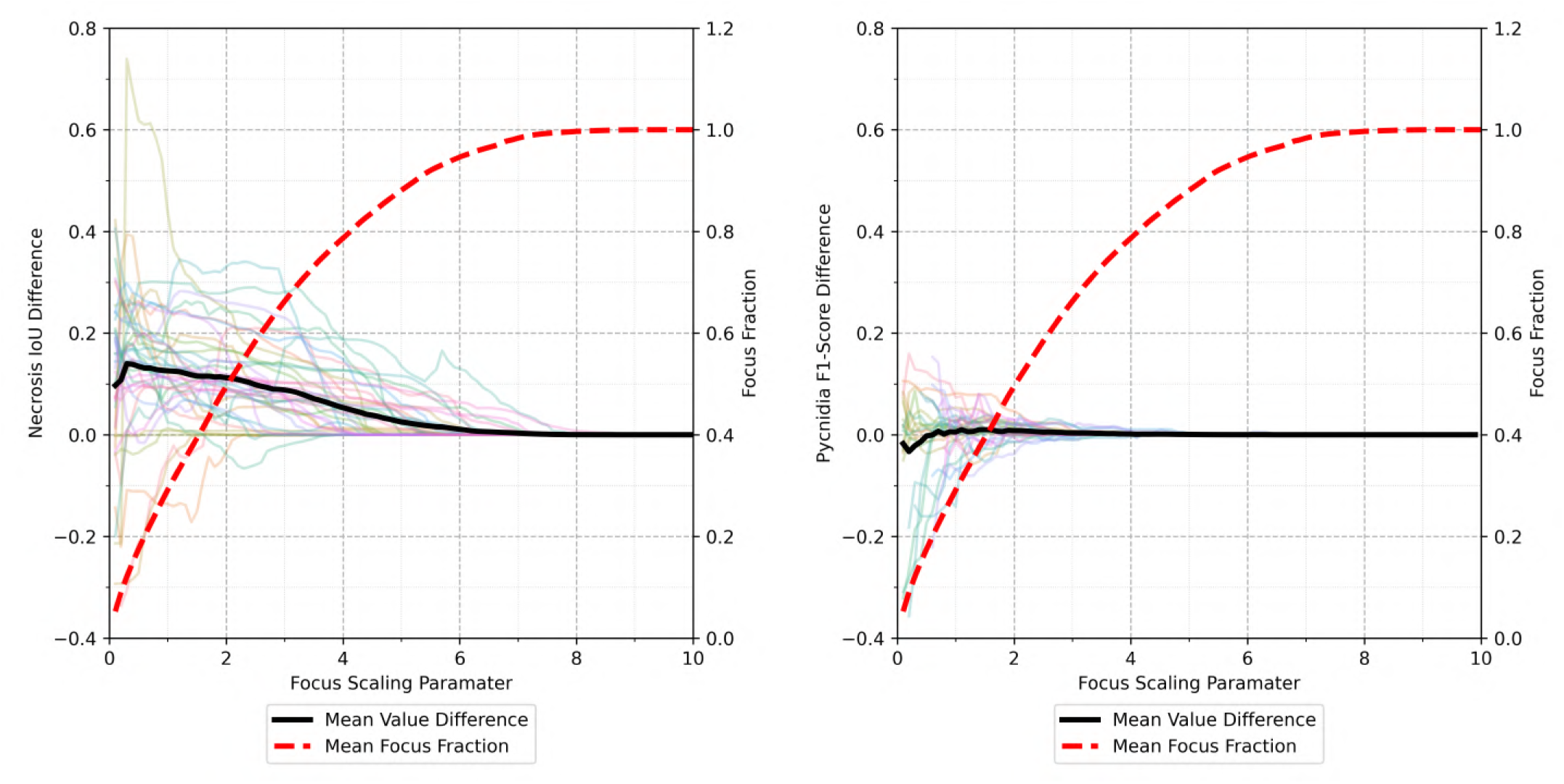
Necrosis IoU and pycnidia F1 score on LIAC dataset in relation to varying the focus estimation parameter *λ*. Individual colored lines show the performance progression on individual images as the focus estimation parameter is adjusted.

Based on this experiment, a reasonable value for the focus parameter (see Section 3.3) was determined to be *λ* = 1 where on average one third of the image was accepted to be in-focus and the necrosis IoU increased on average by approximately 0.125, whereas the impact on the pycnidia performance was negligible.

### 3.3 Plot Evaluation

To provide a link to established phenotyping protocols, the proposed method with the focus estimation parameter determined above was applied to images from the same plots which had been visually scored for incidence and assessed for conditional severity using flatbed scanners. The PLACL values obtained from both the proposed and traditional methods showed a strong correlation (Pearson *R* = 0.83; Figure 6). However, estimated PLACL values were consistently higher for the established than the newly proposed method.

**Figure 6:**
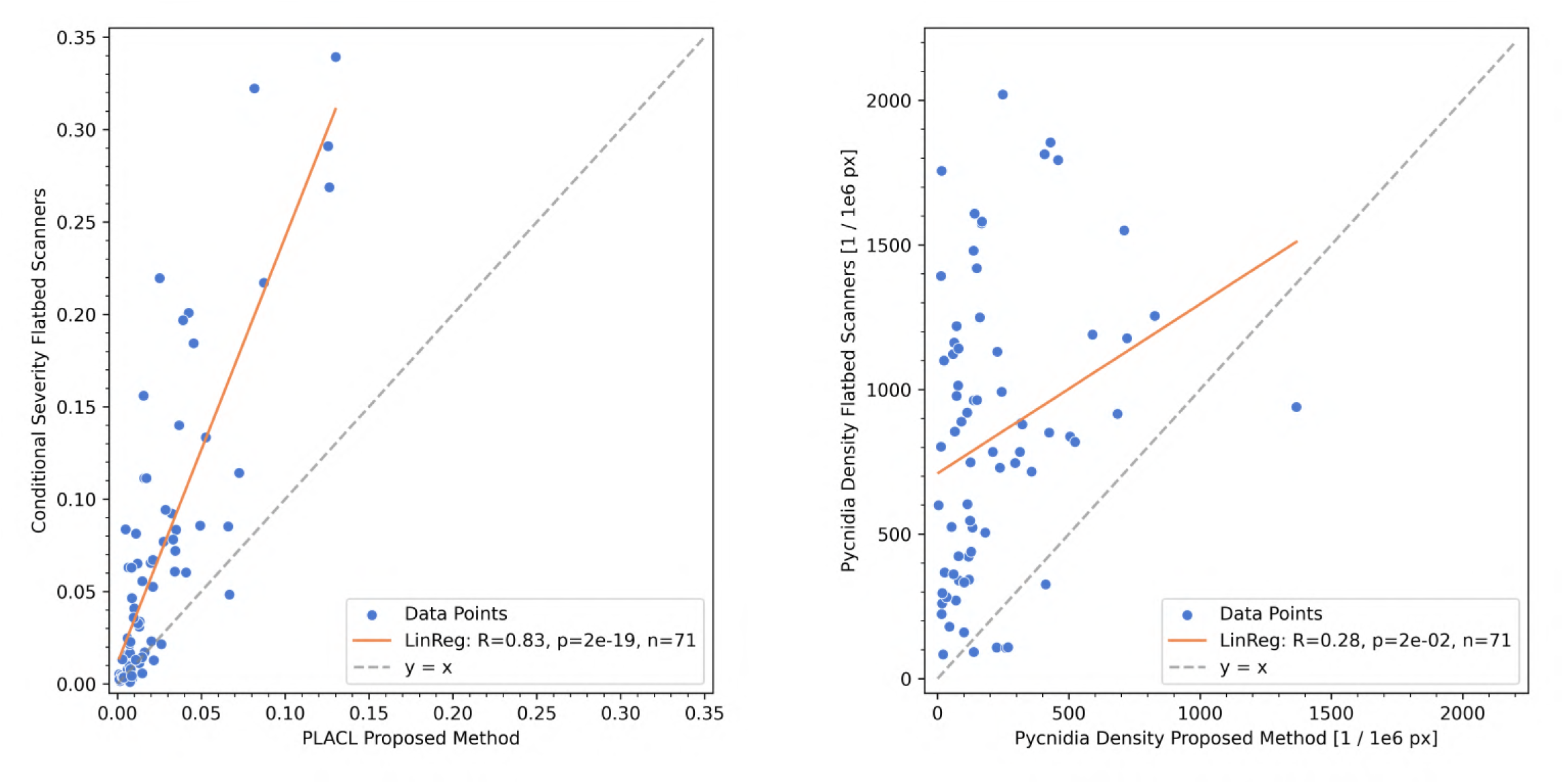
Correlation between estimated values of PLACL and pycnidia density derived from the proposed method and from leaf scans. Pycnidia density is computed with respect to necrotic area.

Pycnidia density correlated weakly (Pearson *R* = 0.28), with the proposed method reporting consistently lower values. Particularly, the proposed method reported lower pycnidia density compared to the flatbed scanner method.

## 4 Discussion

Here, we present a complete workflow for in-field image-based assessment of STB severity. The proposed method directly provides estimates of disease severity from high-quality canopy images. This is achieved by combining organ segmentation, and depth and focus estimation for reference area identification on the one hand with disease detection and symptom segmentation on the other hand. The development of the proposed method relied on newly created training data sets with precise multi-class annotations. Obtained disease severity estimates at the level of field plots were highly correlated with estimates derived from an established disease severity assessment protocol based on time-consuming visual scoring and scanning of sampled leaves [64],[2]. In contrast to these methods or more widely used visual scorings, the data acquisition protocol is based on an approximate point-and-shoot approach, making it amenable to deployment on autonomous machinery to enable fully automated data acquisition.

The core element of the proposed method consists in the segmentation and detection of symptoms in canopy images. For images of unstructured canopy (EFD v2 and EFD v2+) *F* 1*_pycnidia_* decreased from 0.75 [79]to 0.65 (see Table 3) and *IoU_necrosis_* decreased from 0.76 [79] to 0.74 (see Table 2). The decrease in performance was more pronounced for pycnidia than for necrosis (Δ*F* 1*_pycnidia_* = −13.33% Δ*IoU_necrosis_* = −2.6%). We attribute this primarily to the increased complexity of canopy data (e.g., heterogeneous lighting, out-of-focus areas, organs, object orientation in space) compared to flattened leaves, leading to lower confidence of annotations and model predictions due to the increased difficulty of the task. The larger performance degradation for pycnidia detection than for symptom segmentation may be explained by focus blur affecting small-scale features such as pycnidia more than larger objects such as necrotic lesions.

Using only the newly annotated canopy-level data (EFD v2, 400 images) yielded a dataset size similar to the original EFD dataset (422 images) [79]. Training on both canopy images and images of flattened leaves more than doubled the size of the dataset, but resulted in only a slight improvement in symptom detection and a minor drop in segmentation performance (see Tables 3 and 2). Notably, training solely on flattened leaves (EFD) yielded decent but substantially lower performance. This indicated that for dataset extensions, focusing on a specific niche (e.g. canopy images) is more effective than combining diverse data types in a one-size-fits-all approach.

The LIAC dataset was used to evaluate the performance of symptoms detection and symptoms segmentation on large images with more contextual information, more dominant presence of other organs and large out-of-focus regions (see Section 3.2). The performance for pycnidia on LIAC was similar to the validation set, but the performance of necrosis was substantially lower in the LIAC samples (see Figure 4). These patterns can be explained by the more prominent presence of focus blur in LIAC images. For pycnidia this does not make a difference because in these regions no pycnidia can be detected solely due to blur. Note that due to the fixed focal distance, distant regions that would already lack sufficient resolution for pycnidia detection are inherently rendered deeply out of focus. In contrast, for necrosis the image quality cutoff is much more fluid as necrosis can still be annotated and predicted under a significantly higher degree of blur, but with lower confidence and consistency. This aligns with the observed impact of focus estimation on the method’s performance (see Figure 5). Masking highly out-of-focus regions effectively bridged the necrosis segmentation performance gap between the LIAC dataset and the validation set, without impacting pycnidia detection, as no pycnidia were present in the masked areas.

Validation on field data from another site demonstrated the robustness of the method towards different operators, sites, and cultivars (Figure 4). No consistent bias towards the site where the training set originated was observed. For necrosis the performance was even better for the previously unseen site. This can be explained by differences in lighting, growth stage and severity of infection, the contrast between healthy tissue and lesions, as well as co-infections and exposure to other stresses. The absence of performance degradation on an unseen site indicates that the method is sufficiently robust for adoption by other researchers without modifications.

Since scenarios were observed where the performance was worse without any obvious increase in difficulty upon image inspection (see Appendix A.1), it suggests an out-of-distribution inference i.e., predicting on samples not sufficiently covered in the training dataset. We believe that this is due to the training dataset in its current form not adequately capturing the symptom diversity in the presence of diverse settings (e.g. scene composition, lighting). Increasing the dataset size should improve the diversity coverage. However, we advocate for more sophisticated selection of new samples to be annotated in contrast to selecting samples at random. The data acquisition is efficient and thus unlabeled images are abundant. Strategies such as active learning through e.g. uncertainty or diversity sampling can provide a more efficient way to move forward [27] [55].

The need for large and diverse training data sets to support robust deeplearning-based trait extraction from field imagery is widely recognized [17] [70]. For organ segmentation, the ongoing efforts of the scientific community [70] indicate a great general interest in the ability to segment organs for a number of different downstream tasks. In the context of plant disease symptoms, a joint effort to compile datasets across institutions would accelerate the capture of symptom diversity and support large-scale annotation, offering broader coverage, even if site-specific variation is not currently a major performance bottleneck.

However, in the context of STB, to the best of our knowledge, only two datasets provide pixel-wise lesion annotations: the EFD dataset [79] and the PlantSeg dataset [72]. However, we do not consider the PlantSeg dataset suitable for quantifying STB, as its level of annotation detail is insufficient. We rather see its value and the value of other large plant disease datasets [30] [67] [75] [59] [63] [67] [54] as providing an opportunity to pre-train on plant diseases. Currently, the usual starting point for plant disease applications development are readily available imagenet [19] or COCO [38] trained weights. However, these very large general purpose datasets exhibit a large domain gap towards plant diseases. We hypothesize that an intermediate training on a collection of large, less detailed plant disease datasets might provide a fitting intermediate domain, ultimately yielding better performance and faster convergence.

Typically, the experimental unit in resistance breeding is not a single image, but a field plot. Therefore, we aimed to quantify the performance of our method by comparing our estimates of disease severity with reference plot values obtained from a detailed but highly time-consuming manual assessment protocol [64] [2]. It is important to note that estimating the performance of any novel phenotyping method at plot scale is challenging, because obtaining a true reference value may not be possible.

A high correlation between the proposed method and reference method was found for PLACL, indicating the capability of estimating PLACL directly in the field. At the same time, a clear bias was observed with the reference method consistently yielding higher PLACL estimates than the proposed approach (Figure 6). While no definitive explanation for this behavior could be derived from the data, several possible factors were hypothesized. Despite the efforts of the reference method to yield objective results, human operators may introduce bias when selecting leaves, unconsciously favoring those with visible symptoms over a truly random sample (see Section 3.6). Second, the proposed method was limited to imaging from the microplot border, primarily capturing the two outermost rows. Since plot borders often differ in microclimate (e.g. receiving more sun and wind) this could result in differing STB severity, due to factors known to influence the development of STB such as leaf temperature and leaf moisture [14] [29]. Lastly, the time lag of three days between in-field imaging and flatbed scanner imaging may have allowed additional pathogen growth, increasing STB severity.

Besides the individual components of STB like necrosis and pycnidia number, combined measures were also compared. Estimates of pycnidia density in lesions correlated weakly between the two methods (see Figure 6). Given that PLACL showed a much stronger correlation, the weak correlation for pycnidia density in lesions likely originates from challenges in reliably detecting pycnidia and accurately identifying the correct reference area, ensuring the reference area contains only regions where pycnidia can be detected. Since reliable pycnidia detection requires considerably higher image quality than lesion segmentation, it can lead to scenarios where the lesion is correctly predicted but the pycnidia cannot be distinguished in the same image region, leading to a true negative from the point of view of the image but a false negative from the point of view of the true underlying symptoms. In addition, as is the case for PLACL, pycnidia density may be affected by plot border effects and the time lag between the reference assessment and in-field imaging (for more details see above). However the effect of these factors on pycnidia formation is not well studied.

A surprising finding was the performance of Depth Anything v2 model on our data (see Figure 3). Without any need for adjustments, the model yielded very plausible depth estimates of highly unstructured images of a crop canopy, a very task specific environment. Furthermore, the contours of the predicted depth exhibit very sharp and well localized contours which could be used for refinement of predictions of other models e.g. organ segmentation to boost their performance. However, with the current setting, even Depth Anything v2 struggles with partially blurred awns, but note that the current depth estimation pipeline uses quarter resolution, so when high-precision predictions are desired the full resolution could be used. Nonetheless, this underlines the necessity for high resolution imaging when dealing with organ or symptom level evaluation in order to accurately distinguish the relevant regions.

The use of high-frequency features for identification of the focal plane demonstrated compelling results by extracting regions in focus throughout a wide range of scene compositions and imaging conditions. However, the presented approach still contains several shortcomings. First, the approach assumes a uni-modal distribution of features with respect to the depth at which they occur so the focal plane can be estimated by finding the corresponding peak in the features’ depth histogram. But this assumption may be broken depending on the composition of the scene. When there are fewer, or no objects imaged at the focal plane, the features will rather follow a bi-modal distribution, leading to a false estimate of the focal plane. Furthermore, the same effect can arise under the influence of motion blur, when e.g. a leaf at the focal plane moved, decreasing the image quality in that region. Even if the focal plane is estimated correctly, a region suffering from motion blur might lead to issues with tasks critical to image quality, such as incorrect recognition of the reference area affecting pycnidia detection. To reduce motion blur, increasing the aperture is not possible due to depth of field constraints, and raising the ISO is unfeasible due to the resulting increase in noise. The only remaining solution is additional artificial lighting, which was not pursued due to the complexity of its implementation, particularly because of the canopy’s structure, which would create challenging shadow patterns. Assuming an average diameter of pycnidia of 0.1 mm, with 1/60s exposure time mentioned above such distance will be traveled already when the canopy moves at a speed of 6 mm/s. However, estimating the practical allowed motion of the canopy is difficult, since even under the influence of motion blur, there is still high contrast in the direction perpendicular to the motion. Our data was collected on typical days without explicitly considering wind conditions. We did not encounter particularly windy days during the data acquisition. Although the site where INRAE collected the data is particularly favorable for wind due to its proximity to sea, the method demonstrates robustness against the encountered wind conditions. To the best of our knowledge the occasional wind gusts did not cause any systematic motion blur problems in our data. Nonetheless, a direct assessment of image quality would yield more robust results as it operates on the final image quality regardless of which effect influenced it.

While depth estimation from various sensor types has been widely used in the context of agriculture for tasks like autonomy, harvesting, and plant phenotyping [73], it has not been used in the context of de-focus. We believe this work to be the first of its kind to image in-field with a fraction-of-a-millimeter precision which led to a particularly intensive effect of de-focus. We believe that the proposed imaging methodology combined with the focus estimation step, can be useful to other disciplines that require very high imaging precision, such as organ-level phenotyping or detection of small pests, and thus providing a new basis for development of novel applications.

We believe that the greatest challenge of the proposed method in terms of performance is the fast degradation of image quality due to de-focus which greatly contributes to symptoms ambiguity. To mitigate this, we employ the focus estimation step which currently discards 70% of image area on average, decreasing the methods throughput. Fortunately, due to the ease of data acquisition, the decrease in throughput per image can be compensated by collecting more data. However, the symptoms ambiguity negatively impacts both the annotation process [79] and inference since noisy labels negatively impacts trained model performance [34].

Furthermore, the main open question requiring further investigation is the aggregation of a plot value from images. The established visual scoring methodologies typically observe a specific leaf layer such as flag leaves. However, in a realistic canopy these leaf layers tend to overlap, which is then represented in the images of a canopy. Thus, even though the canopy images were approximately aligned with the flag leaves, it does not guarantee only flag leaves are in the images. Particularly in the case of STB which climbs up the plant using water splash dispersal of secondary inoculum, the canopy shows a strong vertical gradient of disease severity in addition to the senescence of lower leaf layers.

Moreover, the height of the camera is adjusted to match the approximate flag leaves height of the imaged cultivar, however when deployed on a moving platform, the height needs to be selected prior to the imaging as the camera is most probably going to be fixed at a constant height. Here we envision a sensor array covering a whole interval of potential heights of interest depending on the underlying cultivar selection and extracting the plant height post image acquisition based on cues like position of wheat ears in the images. Plant height can then be incorporated into the disease assessment to as it is a factor influencing STB severity [62] [61].

From a broader perspective, the method gets rid of the need for any prerequisite knowledge in the context of plant diseases and can be used by any operator with a minimal training. It does not require any specialized equipment. Furthermore, it provides additional insights in the form of symptom level assessment of multiple diseases and insect damage, otherwise not available in the typical breeding workflows, where a single visual assessment of STB severity is done. Even though this work presents an important step forward in terms of usability and throughput compared to previous work [79], it does not solve the most critical prerequisite for its adoption for resistance breeding at scale. Compared to visual scoring, the proposed method still takes longer per plot, making it best suited for deployment on an autonomous ground vehicle (AGV) to enable practical use in commercial breeding. The pipeline requires no canopy interaction, allowing for contact-free imaging with only an approximate camera position at the microplot border. Already the currently used RTK-GNNS enabled agricultural machinery achieves a centimeter-level positioning accuracy, well sufficient for the described tasks. Furthermore, the precision and availability of autonomous agricultural machinery is expected to increase in the near future. Utilizing AGVs as sensor carriers would offer a wide range of benefits in terms of data availability and consistency. Not only will the data acquisition become arbitrarily scalable in terms of experiment size but it will also offer the potential for increased temporal and spatial resolution. Currently, disease severity assessments are often limited to a few or even a single timepoints [4] due to the linked operational efforts. In contrast, repeated assessments increase precision by reducing the impact of errors associated with single measurements. This approach is utilized in the Area Under the Disease Progress Curve (AUDPC), which has been widely used in research, but its application at scale is limited by the throughput of the available disease assessment methods. Repeated assessments also enable tracking disease progression, which is fundamental to understanding interactions between host, pathogen, and environment. Finally, differences in disease dynamics across cultivars can reveal the ability of hosts to delay or decrease the build-up of secondary inoculum [52] This enables the exploration of additional resistance mechanisms not detectable from a single measurement, for example measuring the latent period of STB through artificial inoculation and repeated assessments of the disease to find the timepoint when pycnidia start to emerge.

## 5 Conclusion

In this work, we introduced a novel non-invasive multi-task methodology for in-canopy imaging and recognition and quantification of STB symptoms. The capability of the method to handle the semantics of the imaged scene renders object pre-selection or accurate placement of the camera by the operator unnecessary. This presents a great opportunity for a deployment on AGVs to collect and evaluate data at scale in a fully automated manner, with the potential to provide data at an unprecedented scale and temporal resolution.

While the method achieves high correlation with the established methods, some challenges remain, particularly in accurately detecting pycnidia under varying image quality conditions. Validation on an independent field experiment confirmed the method’s adaptability to different sites without retraining, reinforcing its practical applicability for broader research use.

Future work will focus on optimizing image processing strategies to maximize the analyzable fraction of individual images, refining plot-level metric aggregation methods, and fostering collaborative efforts to expand diverse training datasets. With continued improvements, this approach has the potential to transform disease phenotyping workflows, paving the way toward high-throughput, in-field STB quantification that is accessible, reliable, and scalable for breeding programs.

## 6 Credit Author Statement

R.Z.: methodology, software, investigation, formal analysis, data curation, writing - original draft, visualization. B.A.M.: funding acquisition, conceptualization, writing - review and editing. A.W.: resources, writing - review and editing. M.U.: investigation, writing - review and editing. C.S.: investigation, writing - review and editing. J.A.: conceptualization, investigation, writing - original draft, supervision, project administration.

## Acknowledgments

We want to thank J. Alassimone for supporting during the preparation of pathogen inoculations. Further, we want to express gratitude to the Group of Crop Science at ETH Zürich, particularly S. Corrado for their expertise in crop husbandry and B. Herzog for their assistance in seed preparation and management. We thank RAGT2n for providing seeds of progenies of the population Winner x KWS Ultim and also companies Experando and Lemaire Deffontaines for carrying out and phenotyping the assay in Landrethun le Nord. We also would like to thank S. Vuillemin, S. Gürkan, K. Gefe, T. Khampo, A. Olsen, L. Schwertner and M. Chassot for assistance with data acquisition and image annotation.

## 7 Data availability

The implementation of the proposed method is available at: https://github. com/RadekZenkl/leaf-toolkit The image datasets used in this study can be downloaded from: https://doi.org/10.5281/zenodo.15310826

## Appendix

### A.1 Prediction Examples on Validation Set of EFDv2

**Figure 7:**
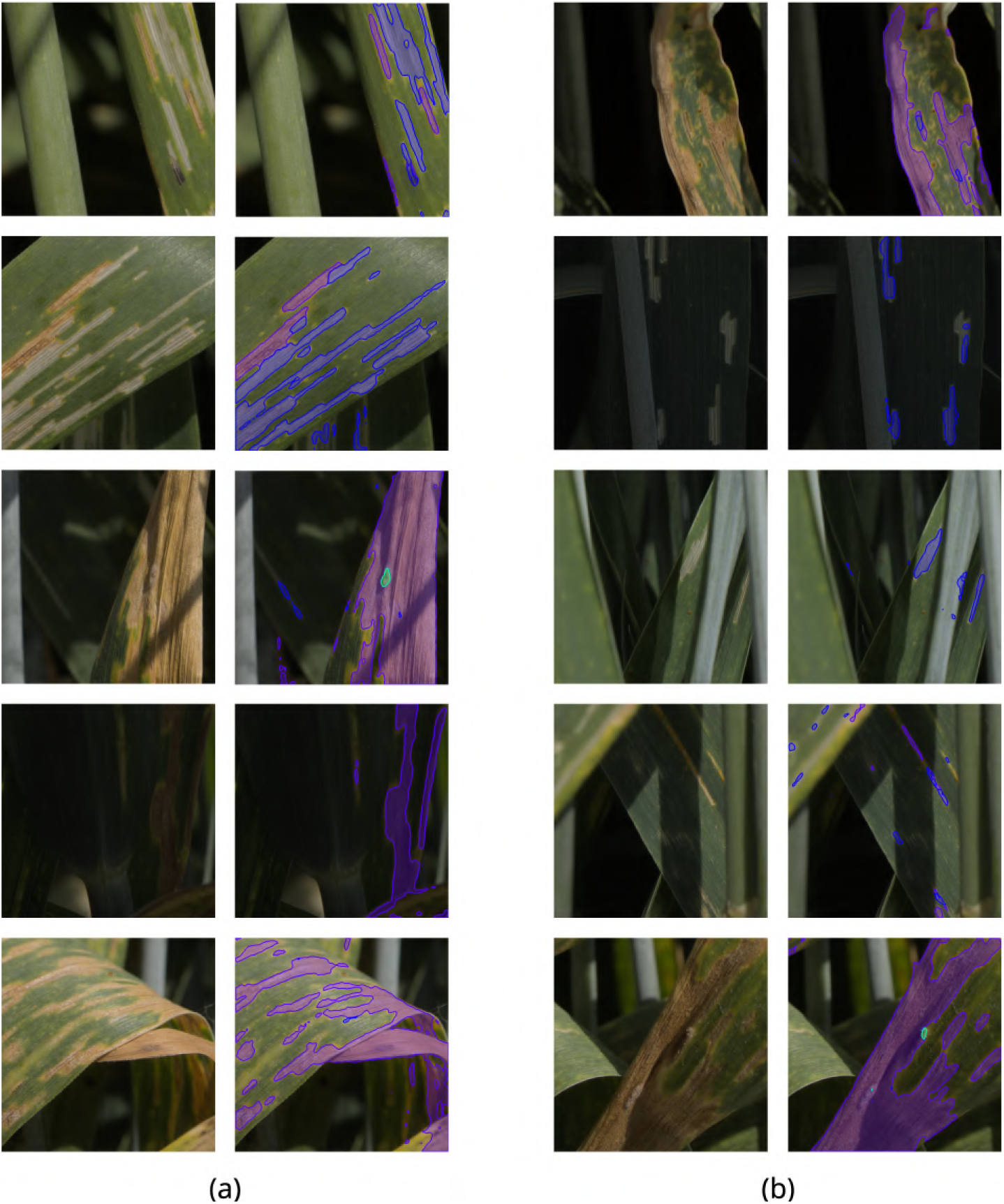
Selected samples from the validation set visualizing success cases panel (a) and failure cases (b). Each image pair contains the input image on the left and visualized predictions on the right. Necrosis is denoted in purple, insect damage in blue and powdery mildew in green.

### A.2 Cultivars selection ETHZ

**Figure 8:**
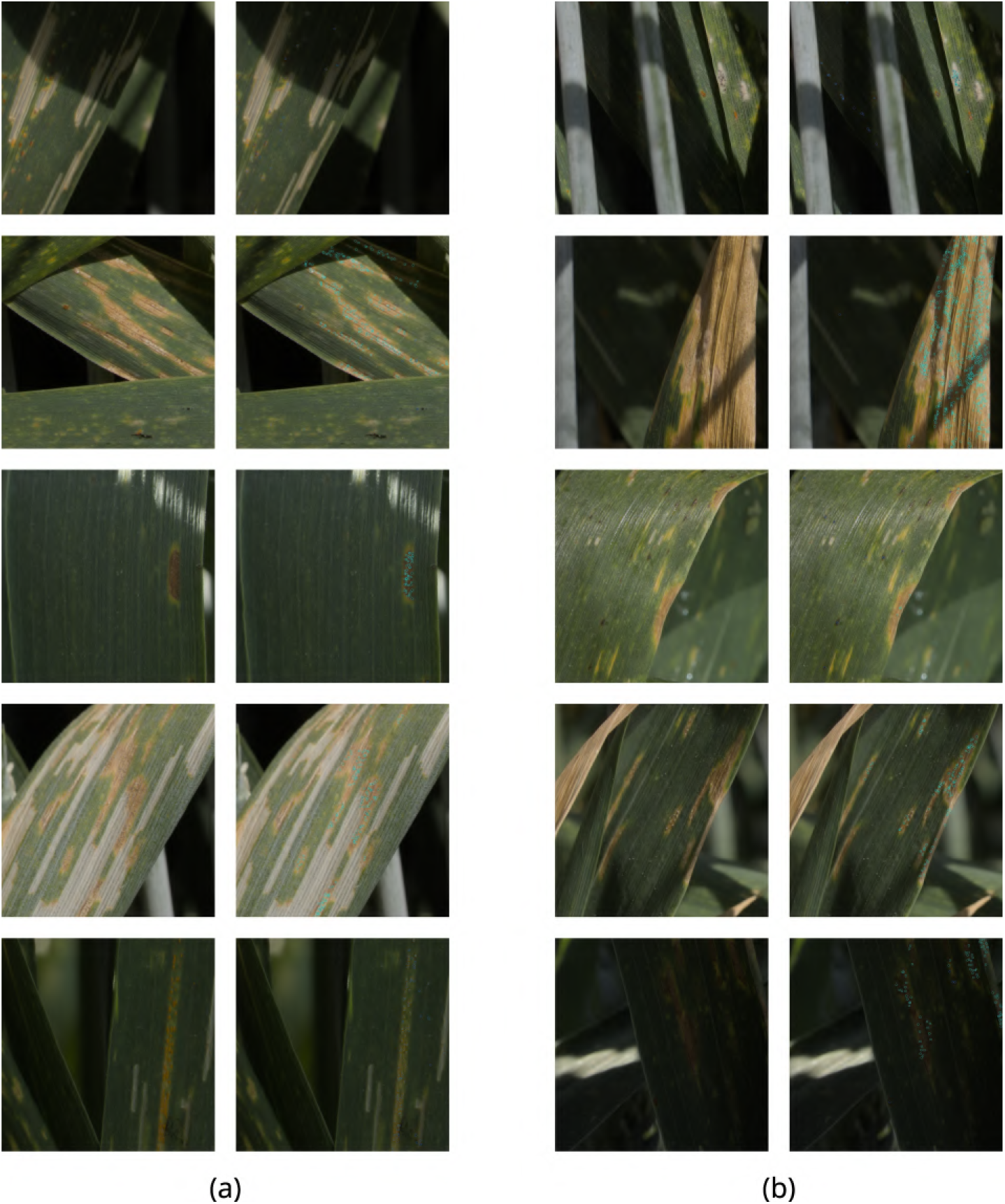
Selected samples from the validation set visualizing success cases panel (a) and failure cases (b). Each image pair contains the input image on the left and visualized predictions on the right. Pycnidia are denoted with cyan circles, rust pustule in blue circles.

**Figure 9:**
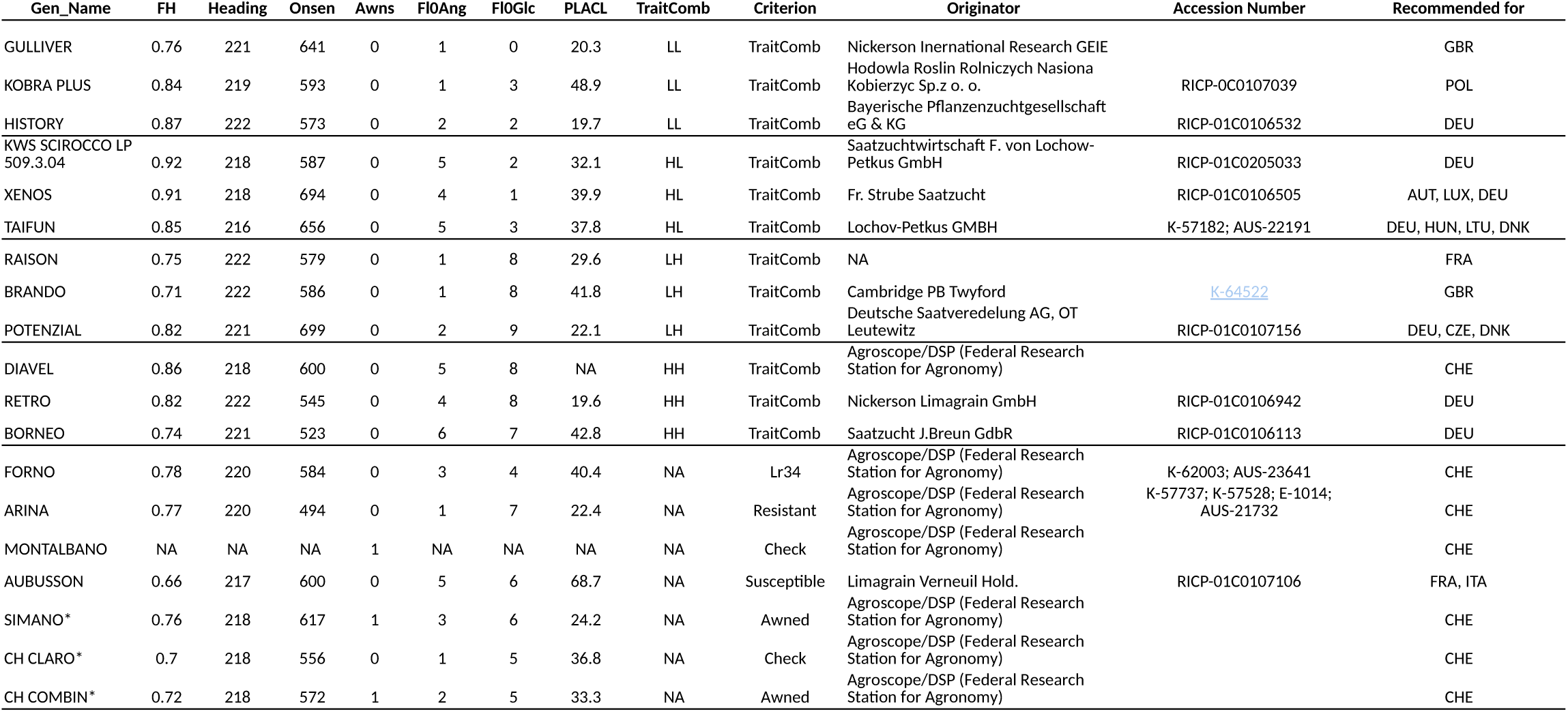
Overview of cultivars from ETHZ field experiment.

### A.3 Plot Imaging

**Figure 10:**
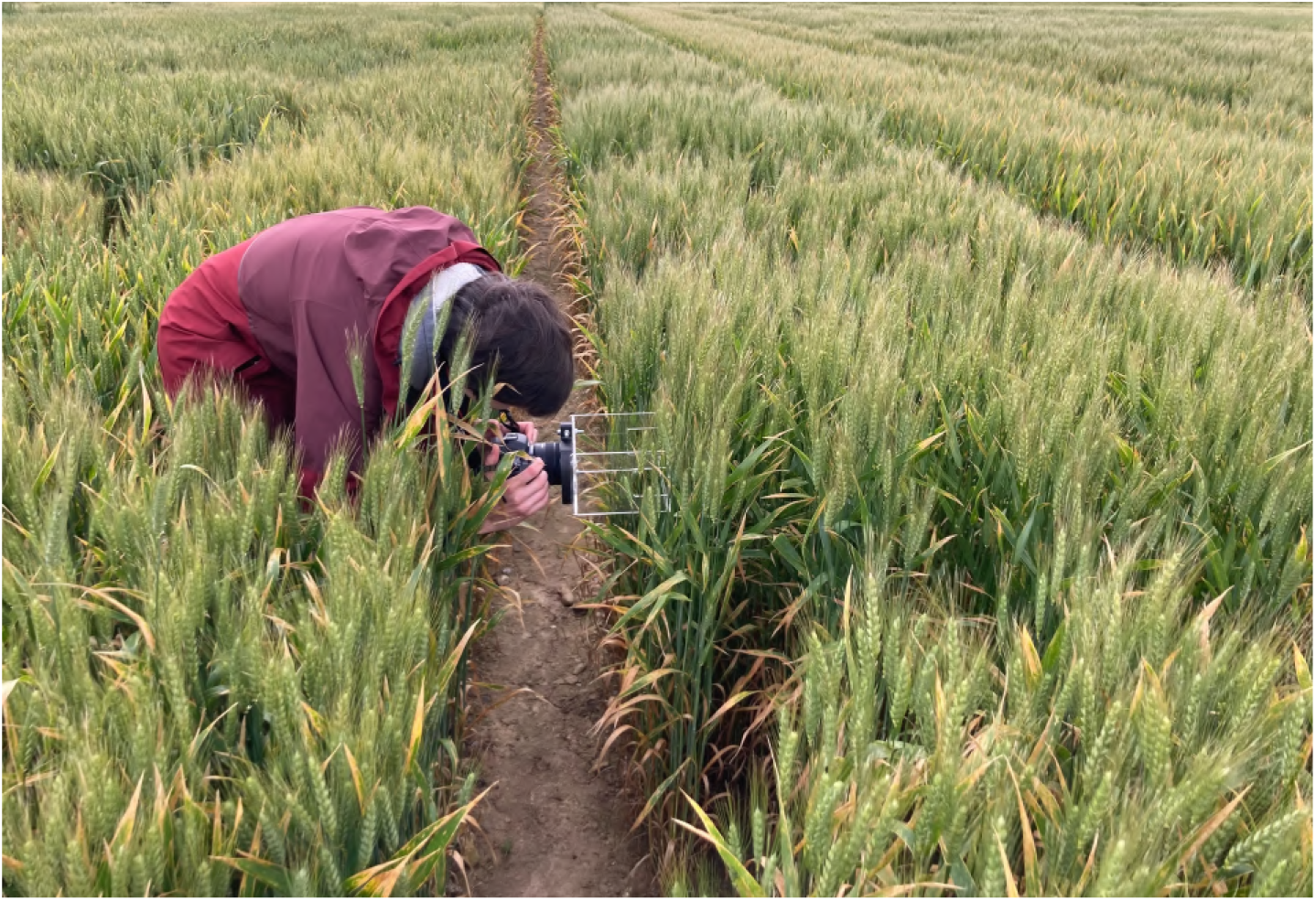
Imaging of a plot along the plot border.

